# Comparative Analysis of Diffusion Models for Enhancing Alzheimer’s Disease Classification

**DOI:** 10.1101/2025.07.25.666665

**Authors:** Jose Gabriel Gonzalez Nunez, Laura J. Brattain

## Abstract

Early and accurate detection of Alzheimer’s disease (AD) is vital for timely intervention and better patient outcomes. However, training machine learning (ML) models for this purpose is challenging due to the limited medical images available and the imbalance of classes. The size and quality of the training dataset directly affect model performance. Recent advances in diffusion models address this limitation by generating synthetic images from a small sample of real images. In this work, we adapted two diffusion models and trained VGG16 and ConvNeXt classification models for AD classification. The first diffusion model was a Denoising Diffusion Probabilistic Model (DDPM) with a U-Net architecture, and the second was a U-KAN framework that integrates Kolmogorov–Arnold Networks (KANs) with the U-Net. Both models were fine-tuned to generate MRI scans of AD or Late Mild Cognitive Impairment (LMCI). We conducted a comparative analysis to assess the reliability and usefulness of these synthetic images for training classification models. The best metrics achieved by the classification models using synthetic images for the AD class were precision of 96%, recall of 83%, F1 score of 87%, and AUC of 0.88. For the LMCI class, the best values were precision of 78%, recall of 88%, F1 score of 82%, and AUC of 0.88. They both demonstrated noticeable improvement from the baseline trained only on the original images.

## 1. Introduction

Alzheimer’s disease (AD) is a progressive neurodegenerative disorder that affects millions of people worldwide, with its prevalence increasing as populations age. Early and accurate classification is essential for effective intervention and management of symptoms. AI models using medical imaging and cognitive data have become valuable tools for classifying MRI images into five severity levels: Cognitively Normal (CN), Subjective Memory Complaint (SMC), Early Mild Cognitive Impairment (EMCI), Late Mild Cognitive Impairment (LMCI), and AD. However, class imbalance poses a major challenge. Generative models like diffusion models offer promising solutions by producing high-quality images from limited data and reducing dependency on scarce real data. [1], [2]

We adapted two diffusion models: DDPM [1] and U-KAN [3]to enhance classification performance using VGG16 and ConvNeXt architectures. The diffusion models were trained to generate MRI images for AD severity levels with limited data. We assess the reliability and validity of these synthetic images when used to augment training datasets for AD severity classification. The implementation process for this study is shown in Figure 1.

**Fig. 1.**
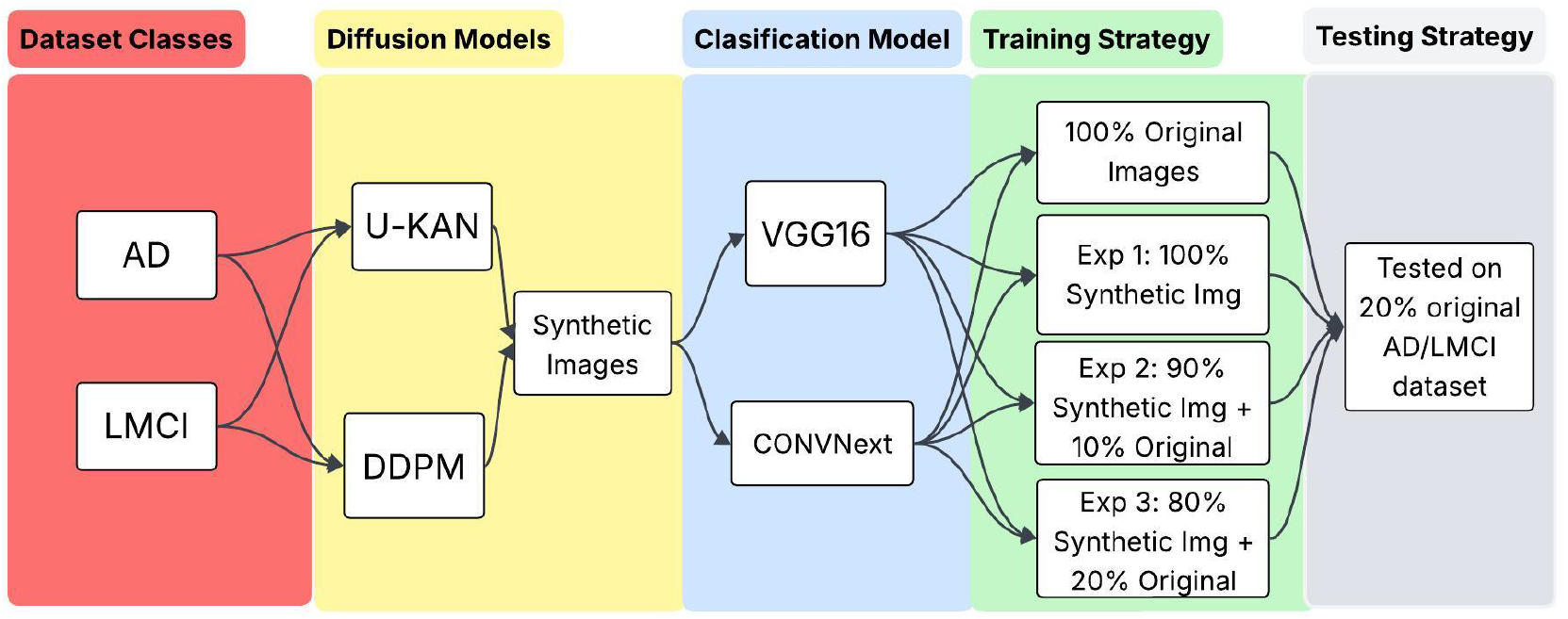
Overview of the proposed experimental pipeline. The diagram shows how AD and LMCI classes are used to train two diffusion models (U-KAN and DDPM) to generate synthetic images, which are then used to train two classification models (VGG16 and ConvNeXt). Four training strategies combine different proportions of synthetic and original images to assess the benefit of synthetic data for classification. All models are tested on a 20% original images dataset.

## II. Related Work

Different synthetic image generators have been developed over the years, with the most commonly used being Generative Adversarial Networks (GANs), Variational Autoencoders (VAEs), and Diffusion Models. GANs are unsupervised generative models composed of a generator that creates data and a discriminator that distinguishes between real and synthetic data. VAEs generate data by encoding inputs into a latent space and decoding them back. Diffusion models consist of two phases: a forward phase that adds Gaussian noise to the data, and a denoising phase that gradually removes the noise to recover the original data.

GANs can generate high-quality images but suffer from unstable training and require careful tuning. VAEs are easier to train on small datasets but often produce less realistic outputs. In contrast, diffusion models generate detailed images, have stable training, and can perform well with limited data through augmentation.

Diffusion models, which add and remove noise in steps, have shown strong performance in medical imaging applications, including image generation and anomaly detection [2], [4], [5].

Deep learning techniques, especially U-Net-based architectures, have significantly advanced medical image segmentation. U-Net is widely used in image generation and segmentation tasks [3]. Recent advances involve modifying U-Net for diffusion models, particularly for segmentation. U-KAN, which integrates Kolmogorov–Arnold Networks (KANs) into U-Net, improves interpretability and efficiency, outperforming other models in both accuracy and speed [3].

The combination of U-Net-based diffusion models with architectures like U-KAN has shown promising results in medical image analysis, enhancing segmentation and anomaly detection tasks [3], [4]. These developments show that diffusion models are key to addressing data scarcity and privacy concerns in medical imaging.

## III. Materials and Methods

### A. Dataset Description

The dataset used in this study was obtained from the Kaggle open source Alzheimer’s Disease dataset [6] and consists of a total of 1,286 brain MRIs. These are classified into five classes, ordered by level of severity: Alzheimer’s Disease (AD), Late Mild Cognitive Impairment (LMCI), Early Mild Cognitive Impairment (EMCI), Mild Cognitive Impairment (MCI), and Cognitively Normal (CN). Images in each class are: AD (171), LMCI (72), EMCI (240), MCI (233), and CN (580). There is a noticeable class imbalance. In this study, we focused on applying diffusion models to the two classes with the fewest images, AD and LMCI.

### B. Classification Model

VGG16 is a CNN (Convolutional Neural Network) that uses blocks composed of multiple 3×3 convolutional layers. Max-pooling layers are interspersed to halve the activation map size. The model ends with two dense layers of 4096 neurons and an output layer of 1000 neurons. The “16” in VGG16 refers to the number of layers with weights: 13 convolutional layers and 3 dense layers, out of a total of 21 layers, including 5 Max Pooling layers.

ConvNeXt is a family of pure ConvNet models designed to rival Vision Transformers in accuracy, scalability, and robustness. Built solely from standard ConvNet components, ConvNeXt introduces modern elements inspired by Transformers, such as adjusted stage compute ratios, a “patchify” stem, inverted bottlenecks, large kernels, and fewer activation/normalization layers. These updates allow ConvNeXt to scale comparably to hierarchical vision Transformers while retaining the simplicity of ConvNets. [7]

### C. Diffusion Models

A diffusion model is an advanced technique in the field of machine learning and multimedia content generation that is used to create images and visual content from input data, usually in the form of random noise. The distinguishing feature of diffusion models is their gradual and controlled approach to transform this initial noise into a final image. Diffusion models use fundamental concepts of probability and statistics to calculate how the noise must change at each step to obtain an image that is consistent and realistic. In this study, we adapted two diffusion models to the AD classification. The first model followed the denoising diffusion probabilistic model (DDPM) framework introduced by [1]. The second model extended a medical-focused diffusion model based on the U-KAN backbone architecture [3], which was specifically designed for medical image segmentation and generation tasks.

### 1) DDPM-based Diffusion Model

Denoising Diffusion Probabilistic Models (DDPMs) are generative models that synthesize high-quality images by reversing a sequence of noisy transformations. In the forward process, a Markov chain adds Gaussian noise until the image becomes nearly random. The reverse process removes the noise step-by-step by training a neural network to predict and denoise each state. DDPMs use a score-matching objective during training to improve noise prediction, resulting in accurate, high-quality outputs.

The U-Net architecture is commonly used in the reverse process of DDPMs due to its encoder-decoder structure, which captures both fine details and global context. The encoder extracts high-level features by reducing spatial resolution, while the decoder reconstructs the image. This design enables progressive denoising, producing results that closely match the original data distribution and perform well on benchmark datasets.

### 2) U-KAN-based Diffusion Model

The U-KAN model integrates Kolmogorov–Arnold Networks (KANs) into a modified U-Net framework for medical imaging. By replacing standard convolutional layers with KAN layers, it captures complex, non-linear patterns more effectively. Based on the Kolmogorov–Arnold theorem, these layers help deconstruct functions and interpret intricate data relationships, enabling U-KAN to outperform traditional U-Net and other segmentation models, especially in detecting subtle diagnostic features.

U-KAN also uses tokenization, dividing feature maps into tokens processed by KAN layers to generate efficient segmentation embeddings. This approach maintains high accuracy with reduced computational cost. Its enhanced feature extraction also improves noise prediction in diffusion models, making U-KAN well-suited for medical image denoising.

### 3) Experiment Strategies

The experiments in this study were conducted using four different setups for each class and each diffusion model, as summarized in Figure 1. Initially, the VGG16 and ConvNeXt classification models were trained and evaluated solely on the original images from the AD and LMCI classes [6], using an 80-20 train-test split. These baseline illustrates the models’ performances without any synthetic images generated by the diffusion models.

For Experiment 1, the classification models were trained using only synthetic images: 150 generated images for the AD class and 100 for the LMCI class from each diffusion model (DDPM and U-KAN). Experiments 2 and 3 then incrementally added real images to the training set, increasing the proportion of original data by 10% in each experiment, while keeping the baseline of 150 AD and 100 LMCI-generated images. This setup enabled a comparative analysis of how the varying mixtures of original and synthetic images from each diffusion model influenced detection performance.

The VGG16 and ConvNeXt models for each experiment were always tested on the same original images that were used for their baseline experiments to preserve the consistency of the results and determine whether the addition of synthetic images to the training process improves the initial performance. We used precision, recall, and F1 score as evaluation metrics for all experiments. For an Alzheimer’s classification model on brain MRI scans, the F1-score was chosen as the primary metric because it balances precision and recall, effectively addressing the critical dual concerns in medical diagnosis. In this specific context, high precision to avoid false alarms and high recall to ensure minimal missed cases are equally important.

## IV. Experimental Results

To evaluate the quality of the synthetic images generated by the diffusion models, we used the following metrics: Fréchet Inception Distance (FID), Inception Score (IS), precision, and recall. These results are shown in Table 1, where it can be seen that the UKAN-generated images achieved higher quality across all metrics. Figure 2 shows a comparison of the images generated by the diffusion models and the original images.

**TABLE 1.** Image Quality Metrics for DDPM and U-KAN on AD and LMCI Classes.

| Model | FID (PCA) ↓ | IS ↑ | Precision ↑ | Recall ↑ |
| --- | --- | --- | --- | --- |
| DDPM AD | 138.3101 | 1.7800 | 0.8600 | 0.2398 |
| U-KAN AD | 131.5280 | 2.0110 | 0.7933 | 0.3099 |
| DDPM LMCI | 127.4667 | 1.9110 | 0.8700 | 0.3472 |
| U-KAN LMCI | 121.3240 | 1.9361 | 0.9000 | 0.3333 |

**Fig. 2.**
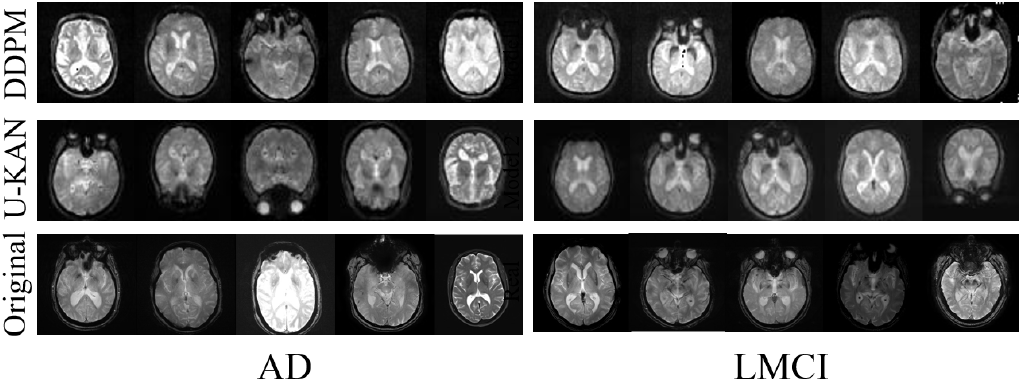
Visual comparison of original and synthetic AD/LMCI MRI scans. Synthetic images from U-KAN and DDPM resemble original data in key features.

The results in Tables 2 and 3, which show the results of the experiments conducted on the VGG16 and ConvNeXt models using the AD class, indicate that a balanced combination of synthetic and real images yields strong performance. DDPM achieves its best result in experiment 3 when the classification models were trained with 20% of real images and 80% generated images. Both classification models showed an improvement of about 15% in precision compared to their baseline models and achieved similar F1 scores. It can also be observed that U-KAN model obtained considerably better results in experiments 2 and 3, obtaining its best scores in experiment number 3 with an F1 score of 87% for the VGG16 model and a score of 75% for the ConvNeXt model, a noticeable increment over the baseline model.

**TABLE 2.** VGG16: Results Comparison for AD Class.

|  |  |  |  | Precision | Recall | F1 |
| --- | --- | --- | --- | --- | --- | --- |
| AD Original Baseline |  |  |  | 0.76 | 0.82 | 0.79 |
| Expt. | DDPM |  |  | U-KAN |  |  |
|  | Prec. | Recall | F1 | Prec. | Recall | F1 |
| 1 | 0.59 | 0.75 | 0.66 | 0.73 | 0.89 | 0.80 |
| 2 | 0.72 | 0.71 | 0.72 | 0.80 | 0.89 | 0.84 |
| 3 | 0.88 | 0.61 | **0.79** | 0.91 | 0.83 | **0.87** |

**TABLE 3.** ConvNeXt: Results Comparison for AD Class.

|  |  |  |  | Precision | Recall | F1 |
| --- | --- | --- | --- | --- | --- | --- |
| AD Original Baseline |  |  |  | 0.73 | 0.65 | 0.69 |
| Expt. | DDPM |  |  | U-KAN |  |  |
|  | Prec. | Recall | F1 | Prec. | Recall | F1 |
| 1 | 0.79 | 0.53 | 0.63 | 0.66 | 0.86 | 0.75 |
| 2 | 0.94 | 0.44 | 0.60 | 0.95 | 0.58 | 0.72 |
| 3 | 0.95 | 0.58 | **0.72** | 0.96 | 0.61 | **0.75** |

Tables 4 and 5 highlight the effectiveness of diffusion models in enhancing performance for classes with limited training data. Initially, the LMCI class was trained with only 58 original images. In later experiments, the training set was expanded to 100 images by incorporating synthetic data, leading to significant performance gains. In experiment 3, the best results obtained with DDPM-generated synthetic images were an F1 score of 71% for the VGG16 model, a 31% improvement over its baseline, and 74% for ConvNeXt, marking a 36% increase. Similarly, with UKAN-generated images, VGG16 achieved an F1 score of 82% (a 42% improvement), while ConvNeXt reached 75%, reflecting a 35% gain over its baseline.

**TABLE 4.** VGG16: Results Comparison for LMCI Class.

|  | Precision |  |  | Recall | F1 |  |
| --- | --- | --- | --- | --- | --- | --- |
| LMCI Original Baseline | 0.45 |  |  | 0.36 | 0.40 |  |
| Expt. | DDPM |  |  | U-KAN |  |  |
|  | Prec. | Recall | F1 | Prec. | Recall | F1 |
| 1 | 0.36 | 0.21 | 0.26 | 0.75 | 0.50 | 0.60 |
| 2 | 0.58 | 0.57 | 0.58 | 0.80 | 0.67 | 0.73 |
| 3 | 0.60 | 0.88 | **0.71** | 0.78 | 0.88 | **0.82** |

**TABLE 5.**
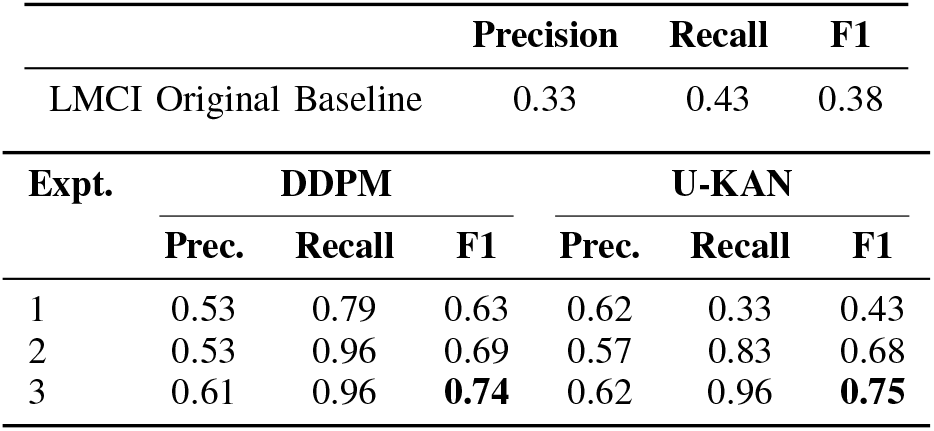
ConvNeXt: Results Comparison for LMCI Class.

A Receiver Operating Characteristic (ROC) curve was also generated to support and illustrate the results shown in the previous tables. Figure 3 shows the ROC of the AD class, including the baseline performance of the ConvNeXt and VGG16 classification models and the best results for each detection-diffusion model combination. Figure 4 presents the corresponding ROC curves for the LMCI class.

**Fig. 3.**
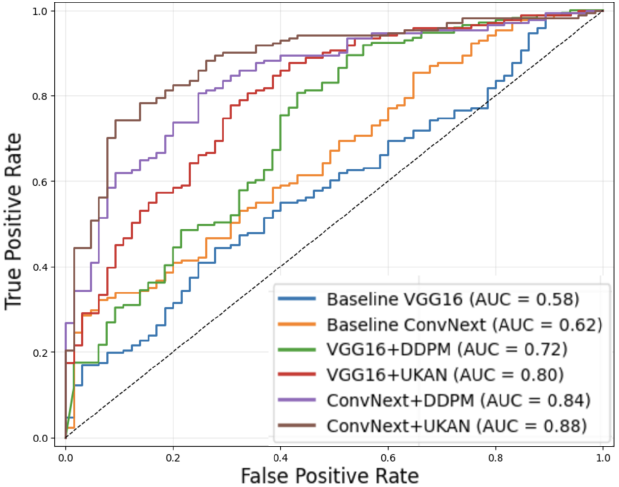
ROC curve for AD class. Using synthetic data improves AUC scores across both classifiers.

**Fig. 4.**
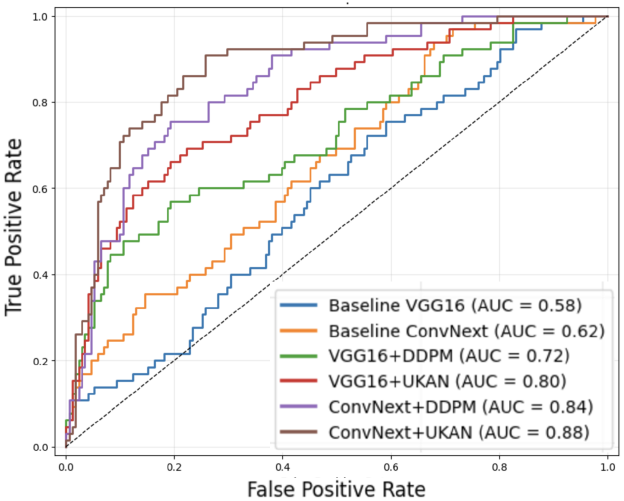
ROC curve for LMCI class. Using synthetic data improves AUC scores across both classifiers.

## V. Conclusion

An accurate classification of cognitive decline is critical in the monitoring and treatment of AD. However, high-severity conditions are typically associated with limited data. To overcome this challenge, in this study, we adapted two diffusion models to generate synthetic brain MRI scans for two minority classes, AD and LMCI. U-Net was integrated with the diffusion models to improve the efficiency and accuracy. A comprehensive comparative analysis was performed to understand the influence of the percentage of synthetic images used in a training set.

Overall, these findings suggest that diffusion models and synthetic images offer a promising approach to address the scarcity of publicly available data in the medical field. By developing specialized diffusion models tailored for medical applications, these technologies could support the training of essential classification models, helping to overcome data limitations.

Future work includes expanding the study to larger AD datasets and other clinical applications that suffer from class imbalance.

## IV. Compliance with ethical standards

This research study was conducted retrospectively using human subject data made available in open access by Kaggle. Ethical approval was not required as confirmed by the license attached with the open access data.

